# Parameterized expertise: Evidence for gradations of musical expertise from electroencephalographic monitoring of auditory decision-making

**DOI:** 10.1101/370379

**Authors:** Jason Samuel Sherwin, Paul Sajda

## Abstract

A musician’s nervous system is thought to specialize to their mode of music production. For instance, the pianist’s control of hands and arms develops to facilitate greater dexterity at the keyboard, while the cellist develops control to play notes using both the fret board and/or bow. Our previous work, employing an anomalous musical event (AME) detection task, identified neural and behavioral correlates that differentiated between a specific class of musicians, cellists, and those without professional musical training and expertise. Here we investigate a fine-grain differentiation between musicians having different modes of musical production, specifically in terms of how these differences are manifested in the neural correlates identified in the AME task. We show, using electroencephalography (EEG), that both event related potentials (ERPs) and single-trial analysis of the EEG can grade musical expertise by mode of sound production. Important is that these fine-grained EEG correlates are observable absent any motor response or active music production by the individuals. We find evidence that these grades of expertise are mediated by different sensory-motor interactions emblematic of the sound production mode. More broadly, our results show that neural markers can both define types of musical expertise and decompose their source components when behavioral differences are either minute or indistinguishable.

## 1 Introduction

Musical expertise has been shown to manifest in brain response across a variety of imaging modalities. For instance, electroencephalography (EEG) experiments have shown that music experts have greater neural response magnitudes in a variety of components in response to unexpected harmonies and pitches (Koelsch et al., 2002b; Koelsch et al., 1999; Tervaniemi et al., 2006; Tervaniemi et al., 1997; Tervaniemi et al., 2005; Tervaniemi et al., 2009). Differences have also been seen using similar paradigms while imaging with magnetoencephalography (MEG) and functional MRI (fMRI) (Doelling and Poeppel, 2015; Schulze et al., 2011). In many cases, these neural differences co-occur with behavioral response advantages seen in musical experts and are linked directly to their behavioral performance gains (Sherwin and Sajda, 2013; Tervaniemi et al., 2005).

An enduring feature that characterizes musical expertise at the neural level is the connection between auditory and motor structures in this class of individuals. Zatorre and others have noted the role of premotor cortex in the timing and organization of actions underlying music production (Zatorre et al., 2007). Others have noted the role of the cerebellum in the production of song (Callan et al., 2007). Maes et al. put forward a theory of music cognition demonstrating the intimate link between music perception and corresponding motor commands (Maes et al., 2014). In particular, they argue that the common coding theory could explain the apparent ease by which sensory perceptions are translated to and from forward and inverse models of movement. Novembre and Keller review behavioral and neurophysiological research in the context of the common coding theory of perception (Novembre and Keller, 2014). They find that this body of evidence leads to musicians’ predictions during both perception and production of music. Drost et al. (Drost et al., 2007) showed evidence for action-effect associations between two groups of instrumentalists (guitarists and pianists). They claim this result reveals a specific component of sensory-motor interaction specific to the instrument specialty. Vuust et al. showed that musical style/genre impacts neural response in musicians (Vuust et al., 2012). Particularly, they found significant differences in response to timbre, pitch and other musical dimensions among jazz, classical, band and non-musicians. This link between auditory and motor structures is likely due to the musicians’ need not only to perceive but also to produce sounds of fine harmonic, melodic, and rhythmic resolution, among other features of sound production. Zatorre and others have noted the links between musical imagery, production and perception at the neural level, differentiating professional musicians from non-professional musicians (Zatorre and Halpern, 2005).

Despite the apparent neural and behavioral manifestations of musical expertise, non-musicians and non-professional musicians learn the rules of musical syntax even without producing music via voice or instrument. They generally build a perceptual sequence expectation that guides their listening experiences. We know this, for instance, because there is a clear neural response when harmonic, melodic or other musical characteristic patterns are violated (e.g., by mistake or otherwise) (Maidhof et al., 2010; Steinbeis et al., 2006). Musical lexicons in novices have particularly been shown to depend on neural structures of the superior temporal sulcus (Koelsch et al., 2000; Peretz et al., 2009).

In earlier work, we reported that non-professional musicians and musical experts (cellists) employ different neural structures in musical sequence expectation. Particularly, we found that motor cortical structure activity was strongest in expert cellists at peak brain response to an anomalous musical event (AME) (Sherwin and Sajda, 2013). Corroborating our results, others have found similar results using slightly different paradigms (Bangert et al., 2006; Haueisen and Knosche, 2001; Hickok et al., 2003). However, the musical expertise literature has largely focused on musicians as a whole, rather than examining the many forms of musical expertise within. Tervaniemi points out this gap and elucidates some general features of different types of musicians on a neural level (Tervaniemi, 2009). In a related point, there has been considerable work on transference of musical ability to the domain of language proficiency (Bidelman et al., 2011; Bidelman et al., 2013; Koelsch et al., 2002a; Koelsch et al., 2004). In such studies, the implicit hypothesis is that neural structures facilitating specialized auditory perception and production skills are mediated by auditory-motor capabilities that are beyond the general population of normal subjects. Lacking however is a particular experiment to illustrate or test the hypothesis that differences in specific music production are revealed at the neural level.

Here we build upon our earlier work where we identified EEG correlates that differentiated musicians (cellists) from non-musicians, (Sherwin and Sajda, 2013). Specifically in this paper we examine the extent to which differences in the EEG, both event related responses (ERPs) and single-trial analysis, expose differences in the mode of music productions across several groups of musicians. Similar to our previous study (Sherwin and Sajda, 2013) we modified a well-known cello piece to create a target detection experiment in which we examined subjects’ responses to anomalous musical events (AMEs) embedded therein. As before, the task was passive and did not have the subjects actively play nor respond during the musical piece. We recruited musicians and established groups based on their specific mode of expertise in music production, relating these to the cellist group. We hypothesized that subjects whose method of sound production expertise were increasingly similar to the cellist would show graded neural and behavioral responses to this AME detection task.

## 2 Materials and Methods

### 2.1 Subjects

Twenty-seven subjects in total participated in the study (18 males, 31.5±2.4 year). There were five groups of at least five subjects each. We defined each group by performance similarity to the cellist mode of sound production (henceforth referred to as “cellist expertise”). In particular, similarity of instrumental/vocal specialization to cello performance determined group membership. For instance, a violinist is more similar to a cellist than is a trumpeter because the violinist plays a string instrument. Complete background details of age, age of music training start, age of voice/instrument training start and handedness are shown in the Supplementary Material as Table S1. The “Cellist” (C) group contained professional cellist subjects with a mean age of 41.2±10.6 years (2 males). Individuals in the C group had begun cello performance training at the mean age of 8.6±0.5 years. The “Non-Cellist String Player” (NCSP) group contained professional violinist subjects had a mean age of 33.2±2.8 years (3 males). The NCSP group had begun violin performance training at the mean age of 6.8±1.0 years. The “Singer” (S) group contained professional singer subjects who had a mean age of 36.7±4.2 years (4 males). The S group had begun voice performance training at the mean age of 7.4±1.6 years. The “Non-String Instrumentalist” (NSI) group contained professional trumpeter subjects who had a mean age of 26.8±3.6 years (5 males). The Non-String Instrumentalist group had begun trumpet performance training at the mean age of 8.6±0.9 years. Finally, the “Non-professional Musicians” (NM) group contained novice subjects who did not have any years of professional instrumental/voice performance training and as a group had a mean age of 30.8±5.8 years (3 males). NM were selected based on their having no prior experience playing the cello, though non-professional musical instrument experience was allowed. All subjects reported normal hearing and no history of neurological problems. Informed consent was obtained from all participants in accordance with the guidelines and approval of the Columbia University Institutional Review Board.

### 2.2 Stimuli Overview

As the musical stimulus, we used the first 65 seconds of Yo Yo Ma’s recording of the Prelude of J.S. Bach’s Cello Suite No. 1 (Ma, 1998), i.e., the first musical section of the piece. Subjects listened to this complete musical section, or trial, 40 times, with 32 of these trials altered to contain 3 to 6 anomalous key-change events. Key-shifts were added to the original 44.1kHz .mp3 file using Apple Inc.’s Logic Express 9.0 (Cupertino, CA). These key-changes were inserted at arbitrary times with Pitch-Shifter, a built-in plug-in to Logic Express, and each trial was then saved as a 44.1kHz .wav file. Neither the frequency, timing, nor direction of key-changes in one alteration could be predicted from those of another. The rule for the key-changes was that the net key-change should be no more than a half step from the original key of the recording.

In the case of the Bach prelude (originally in G), the recorded key was altered no higher than G-sharp (G#) and no lower than G-flat (Gb). This was done to avoid overt distortion of the original recording, thereby making a key-change obvious from non-harmonic considerations. We balanced the direction of key-changes using a total of 69 key-changes up and 71 key-changes down. Considering each key-change as a stimulus, the inter-stimulus-interval (ISI) was at least 6 seconds. It was not necessary for the recording to begin in the original key and so some trials began one semitone down or up from the original key to ensure that the task was specific to relative key-change.

These 8 control and 32 key-change trials were presented to the subjects in a randomized order. An example of full block implementation can be found illustrated in Supplementary Material (Figure S 1).

A Dell Precision 530 Workstation was used to present the audio stimuli with E-Prime 2.0 (Sharpsburg, PA) and a stereo audio card. The subjects sat in an RF-shielded room between two Harmon & Kardon computer speakers (HK695-01, Northridge, CA) connected to the Dell. Each subject was allowed to adjust the volume so that they could comfortably perform the task. A schematic representation of a key-change trial is shown in Figure 1.

**Figure 1.**
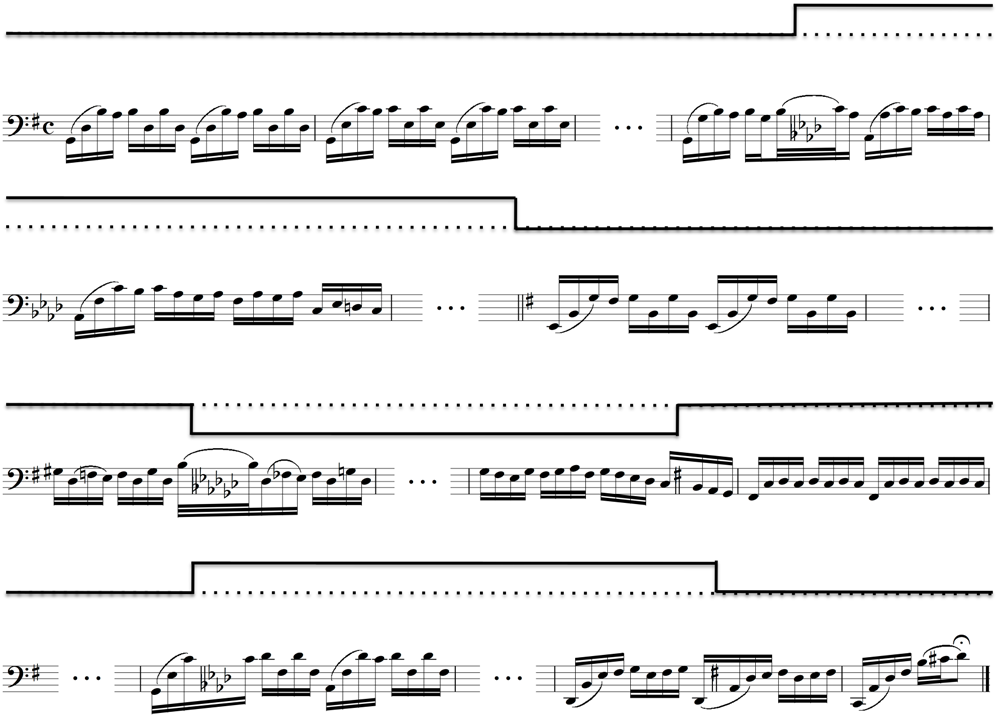
Musical notation and representative step-function of an alternation track showing the semi-tone key-changes imposed on Bach’s *Cello Suite No. 1*, *Prelude*. Above each stave, a step-function shows the key of the music with respect to the original key of G Major. The dotted-line represents the original key of G. Ellipses between bars represent music that remained in the preceding key. It is important to note from this figure that the key-changes were not limited to occur between notes or between musical phrases. Rather, they could occur during notes (e.g., the first and third) or between them, as well as between musical phrases. Although this example begins and ends in the original key, this was not true for the remaining alteration tracks (Adapted from (Sherwin and Sajda, 2013)).

In the figure, a solid step-function above the musical notation provides a schematic of the key for those not fluent in musical notation.

Subjects performed a simple detection task in which they were asked to covertly count the number of AMEs they detected. Counting was used rather than an overt behavioral response, such as a button-press, to minimize motor confounds in the EEG. Counts provided an estimate of task performance and ensured that subjects attended to the task. Subjects were not explicitly instructed to detect a key-change, but simply to attend to anything out of the ordinary – i.e., an anomaly. They were instructed not to move during the task and they were monitored constantly to ensure this. Stimulus events were encoded and synchronized with the EEG recordings via a TTL pulse in the event channel. In post-hoc analysis, stimulus events were added to the EEG of the control tracks.

### 2.3 Data Acquisition

EEG data was acquired in an electrostatically shielded room (ETS-Lindgren, Glendale Heights, IL, USA) using a BioSemi Active Two AD Box ADC-12 (BioSemi, The Netherlands) amplifier from 64 passive scalp electrodes. All channels were referenced to BioSemi’s ground electrodes made for use with the Active Two. Data were sampled at 2048 Hz. A software-based 0.5 Hz high pass filter was used to remove DC drifts and 60 and 120 Hz (harmonic) notch filters were applied to minimize line noise artifacts. These filters were designed to be linear-phase to minimize delay distortions. Stimulus events – i.e., key-changes – were recorded on separate channels.

Throughout the experiment, subjects listened to the music excerpts with their eyes closed. This minimized blink and eye-movement artifacts. This technique has been used in other music perception studies (Maidhof et al., 2010), as well as auditory oddball studies (Goldman et al., 2009). Consequently, no eye calibration experiments were needed before implementing the filtering described above. In epoching the data, the average baseline, from −1000ms to stimulus onset, was removed. After epoching, the ‘poprej’ automatic artifact epoch rejection algorithm from EEGLAB (Delorme and Makeig, 2004) was run to remove all epochs that exceeded a probability threshold of 5 standard deviations from the average.

### 2.4 Behavioral Paradigm and Analysis

At the end of each trial, we asked subjects to report the number of anomalies they detected. Since the AMEs were actually key-changes, we calculated behavioral accuracy by noting the deviation from the actual number of key-changes in each trial. For instance, if excerpt *i* contained *n_i_* key-changes and a subject reported *n_i_*-*k_i_* or *n_i_*+*k_i_* key-changes then this constituted an error of *k_i_*. However, if the subject reported *n_i_* then this constituted an error of zero, *k_i_* = 0. To summarize the performance of each subject, we subtracted from 1.00 (perfect accuracy) the total number of errors, normalizing by the total number of actual key-changes that occurred (Equation 1). Despite the possibility of accuracy being less than zero, this case would only be in the event of extremely poor behavioral performance and did not occur for any subjects in our experiment.

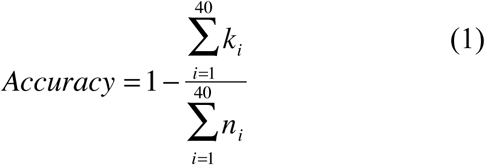

We performed a one-way ANOVA and performed a post hoc correction for multiple group comparisons to examine specific group differences in behavioral accuracy.

We used the Spearman rank coefficient to test linear variation of behavioral accuracy with the hypothesized similarity to cellist expertise. In the absence of any specific guidelines, the Spearman coefficient allowed us to test an ordered variation of behavioral accuracy without specifying *ad hoc* the similarity to cellist expertise.

### 2.5 Traditional Event-Related Potential EEG Analysis

We performed a standard event-related potential (ERP) analysis on the filtered, epoched and artifact-removed (i.e., preprocessed) EEG data. We examined all electrodes by calculating the difference ERP from [−200, +1000] ms. We defined the difference as that between AME epochs (in either direction, regardless of starting and ending key) and control epochs of the same piece of music. To quantify the group differences, we performed a one-way ANOVA on ERPs and performed a post hoc correction for multiple group comparisons in the P300 window (defined as [350, 450]ms). As for behavioral performance, we also used the Spearman rank coefficient again to test linear variation of ERP metrics with similarity to Cellist expertise.

### 2.6 Single-Epoch Classification of Anomalous Musical Events (AMEs)

We performed a single-trial classification of the preprocessed EEG to discriminate between AME and control epochs. This process and its application to the classification of single epochs are illustrated in Figure 2A.

**Figure 2.**
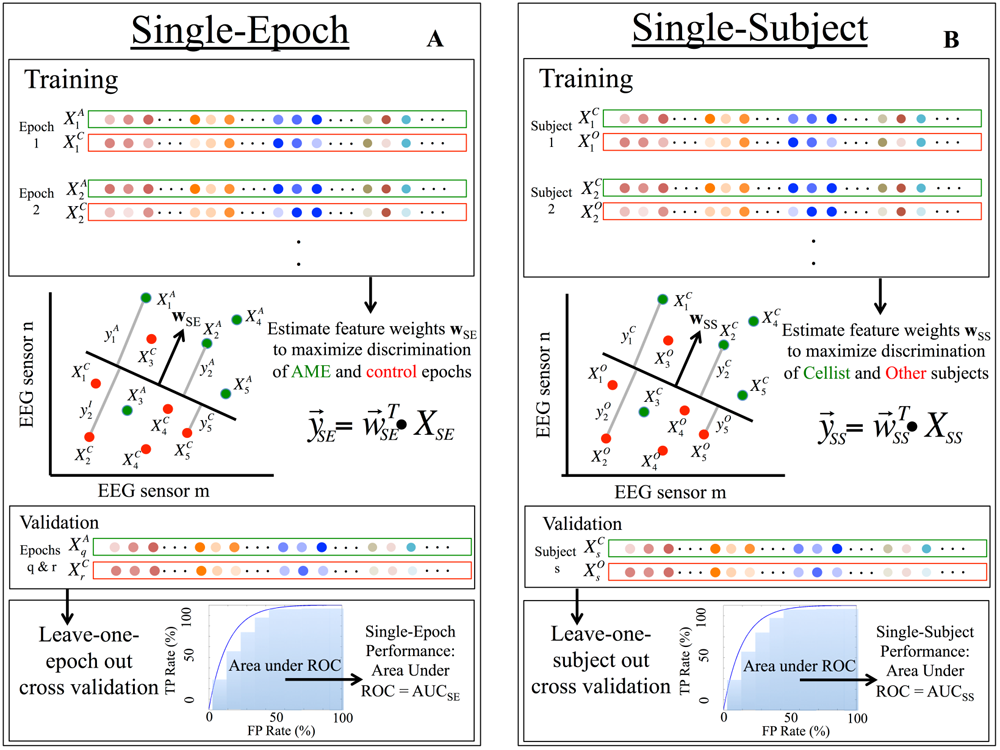
Illustration of single-epoch and single-subject classification training and validation. (Single-epoch classification was done on AME vs. control epochs, while single-subject classification was done on cellist (C) vs. other group memberships (e.g., Non-professional Musicians (NM), Non-String Instrumentalists (NSI), etc.). (**A**) Leave-one-subject-out cross validation of single-epoch and (**B**) single-subject feature vectors. Single-trial EEG data feature vectors (X_SE_) are discriminated by a classifier, **w_SE_**, and trial-averaged difference ERP feature vectors (X_SE_) are discriminated by another classifier, **w_SS_**. Both classifiers are cross-validated with the left-out feature vectors (i.e., the q^th^ or r^th^ epoch for X_SS_, and the s^th^ subject for X_SE_) and the respective Area under the ROC (AUC) metrics are calculated for classification.

We used regularized logistic regression to optimally discriminate between the two epoch-conditions over a given temporal window (Parra et al., 2002; Parra et al., 2005). In particular, we defined training windows (τ) starting at 200ms pre-stimulus and ending at 1000ms post-stimulus, each having a duration of δ = 50ms. The window onset time (τ) was varied across time τ∈[−200,+1000]ms in 25ms steps. We chose this range of τ to provide substantial time both before and after the stimulus to be sensitive to electrophysiological changes that are due to anticipation of the AME. We then used regularized logistic regression to estimate a spatial weighting vector at each value of τ and δ, 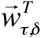, that maximally discriminates between EEG signals X for each condition (e.g., AME versus controls):

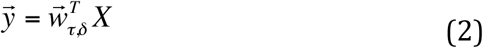

In Equation 2, X is an N x T matrix (N sensors and T time samples). After applying the optimal discrimination vector, 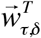, the discriminating component, 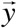, that is maximally correlated with each condition while minimally correlated with both conditions. We use the term ‘component’ instead of ‘source’ to make it clear that 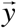 is a projection of all activity correlated with the underlying condition-source. We used the re-weighted least squares algorithm (Jordan and Jacobs, 1994) to learn the optimal discriminating spatial weighting vector 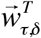, or specifically for epoch-classification, **w**_SE_, and for subject-classification, **w**_SS_.

We quantified the performance of the classifier, 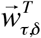, by the area under the receiver operator characteristic (ROC) curve with a leave-one-out approach (Duda, 2001). Throughout the text, we refer to this metric as the area under the ROC, i.e., the AUC. The AUC metric measures discrimination performance in each training window, τ.

We quantified the statistical significance of AUC in each window (τ) with a permutation procedure. We randomized the truth labels between control and AME epochs and then retrained the classifier on these permuted labels. We did this procedure 250 times for each subject. We then constructed a significance line across all windows from these results and Bonferroni corrected for multiple window comparisons (p = 0.05).

We examined the time course of AUC values both between and within groups. To summarize group differences, we examined the window of maximum AUC for each subject group. We used the time and value of maximum AUC as a metric to examine the gradation of neural response seen across the groups. In particular, we performed a one-way ANOVA across groups and followed up with *post hoc* single group comparisons corrected for multiple comparisons. As for behavioral and ERP metrics, we also used the Spearman rank coefficient to test linear variation of single-trial classification metrics with similarity to Cellist expertise.

### 2.7 Single-Subject Classification of Expertise from ERPs

We also trained and tested an elastic net regularized logistic regression classifier to identify group membership on a subject-by-subject basis (see Figure 2B), using leave-one-subject-out per class for cross validation. In particular, following Equation 2, X is now an N’ x T’ matrix (N’ sensors of mean voltage and T’ subjects). We used an elastic net classifier because of the lower relative value of T’ to N’, as opposed to the case for T to N. We utilized an elastic net learning algorithm to solve for the optimal ‘discriminating component’ 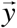 that is specific to activity correlated with each group while minimizing activity correlated with both groups. As before, this algorithm learns an optimal discriminating spatial weighting vector 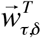, or specifically **w**_SS_.

The use of the elastic net has the effect of reducing potential sources of noise in the discrimination. We implemented the elastic net in our logistic regression (Conroy and Sajda, 2012) by performing a parameter sweep of the sparsity parameter (α) on [0,0.1:0.2:0.5], where α = 0 indicates no sparsity in the solution. When α = 0 (no sparsity) evaluation of the classifier was simply the AUC metric. However when α > 0, we followed the approach of Conroy et al. (Conroy and Sajda, 2012; Conroy et al., 2013) to pick the optimum solution based on evaluating the Pareto frontier—i.e. the convex hull defining the trading off both between the classifier AUC and consistent selection features An example Pareto frontier is shown in the Supplementary Material, Figure S2. In particular, AUC is traded against the mean probability of feature selection (μ_sp_), as defined in Equation 3, where *Ā* is the mean number of features selected by the classifier, p is the number of features and *𝒱_i_* is the i^th^ feature.

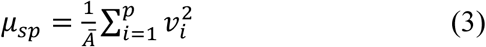

The second parameter (λ) for the elastic net was determined using an algorithm based on degrees of freedom in the solution space (Park and Hastie, 2008). In the Pareto space, the optimum classifier is the one that is that closest to (1,1), indicating an AUC of 1 and mean probability of feature selection of 1. This Pareto optimal criterion was used as the metric for choosing the best classifier for the classification of a subject’s expertise and it is the AUC of this classifier we report in the Results. We calculated p-values via permutations of the subject group labels (10000 permutations). All classification results are reported at p < 0.05.

We examined nine (9) equally spaced post-stimulus temporal windows ([0, 800]ms) for characterizing difference in the ERPs between the groups. As compared to traditional ERP and single-trial analyses, we picked this smaller time period because those results had generally shown a return to baseline by approximately 800ms, with no deflection in general before the AME. Another consideration in the choice of the number of equally spaced windows is that we aimed to minimize the number of multiple comparisons in time, in addition to those across different subject groups. The Bonferroni corrected values of AUC – accounting for both groups (5) and windows (9) – was then calculated as a reference point against which to judge the true AUC values at each window.

### 2.8 Source Localization Analysis

We used low-resolution tomography (sLORETA) of scalp potentials (Pascual-Marqui, 2002; Pascual-Marqui et al., 2002) to identify the cortical generators underlying the differences between groups. To compare groups, we examined differences between AME and control trials. To increase statistical power, we separated key-changes up from key-changes down and computed sLORETA estimates separately. Earlier results showed that there were no significant differences between these anomaly types (Sherwin and Sajda, 2013).

With this increased statistical power we used statistical non-parametric mapping (SnPM) for comparing groups in the sLORETA voxel space. We calculated sLORETA fits for each subject’s grand average ERP at the subject specific maximum AUC for up and down epochs, respectively. The ERP calculations were averaged across time in the 50ms window of peak discrimination between each AME type and controls. We then compared the ten (i.e., five subjects by two AME types) cellist (C) sLORETA fits with the ten corresponding fits of each of the other four groups (NCSP, S, NSI, NM). The sLORETA parameters used in the independent groups t-test for all of the comparisons can be found in the Supplementary Material (Table S6). We established significance using 1024 permutations and the SnPM procedure for voxel-space comparisons (Holmes et al., 1996; Nichols and Holmes, 2002; Pascual-Marqui, 2002; Pascual-Marqui et al., 1999).

## 3 Results

### 3.1 Expertise Graded with Behavioral and Traditional ERP Analysis

The behavioral accuracy and traditional ERP analysis across the five subject groups is shown in Figure 3. Behavioral accuracy as quantified by Equation 1 is shown in Figure 3A. The height of each bar is the mean behavioral accuracy and the error bars indicate standard error about the mean (SEM). Color-coding indicates each group.

**Figure 3.**
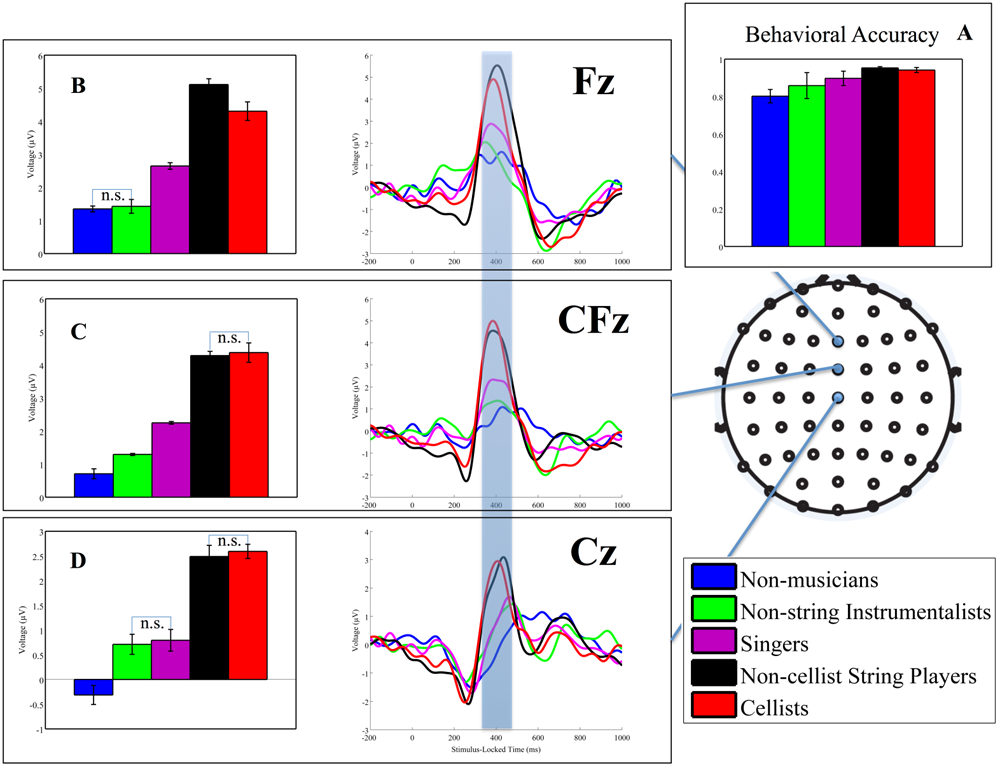
Behavioral results and differences in event-related potentials as a function of subject’s mode of music production. Inset A shows mean behavioral accuracy as defined in the text for each group with standard error of the mean across subjects shown. After correcting for multiple group comparisons, the one-way ANOVA showed no significant differences, though single comparisons between either Non-cellist String Players (NCSP) or Cellists (C) and Non-professional Musicians (NM) showed significant difference (p < 0.05, uncorrected for multiple group comparisons). The 64-channel cap diagram is shown to the right with the three electrodes whose ERPs are shown indicated in blue. The legend indicates group membership: blue = NM, green = NSI (i.e., trumpeters), magenta = S, black = NCSP (i.e., violinists) and red = C. Insets B, C and D show means and standard errors of means across subjects of the blue-shaded time region. After correcting for multiple group comparisons, the one-way ANOVA on group-mean time course showed significant differences; *non*-significant differences are indicated with bars in each inset.

We used a one-way ANOVA to quantify differences between groups for behavioral accuracy. The one-way ANOVA for group difference was significant (p < 0.05, F_(4,24)_ = 3.08). In *post hoc* single group comparisons, corrected for multiple group comparisons with Bonferroni, NM were significantly worse than both Non-Cellist Instrumentalists (p < 0.02, independent groups t-test, t(8) = −3.35), and C (p < 0.01, independent groups t-test, t(8) = −3.83).

Also, we found a general increase in behavioral accuracy with similarity to cellist expertise. From Figure 3A, we see such a trend in the group means as we progress from left to right, i.e., from least to most similarity with cello expertise. We quantified this observation with the Spearman rank coefficient of behavioral accuracy as an assumed linear function of similarity to cello expertise. We found that behavioral accuracy linearly varied with similarity to cello expertise (ρ = 0.54, p < 0.01).

The mean group difference ERPs for the midline electrodes is shown in Figure 3. Color-coding is the same used for behavioral accuracy. The blue shaded bar centered on 400ms indicates the window in which ERP values were compared using a one-way ANOVA. We chose this window because of its co-occurrence with the window of the P300 (Linden, 2005; Picton, 1992). The one-way ANOVA showed significant difference between groups at each of the indicated electrodes (Fz: p << 0.01, F_(4,254)_ = 890.35; FCz: p << 0.01, F_(4,254)_ = 1059.49; Cz: p << 0.01, F_(4,254)_ = 388.55). In *post hoc* single comparison tests, correct for multiple group comparisons, nearly all groups showed differences from each other. Only those differences that were *not* significant are indicated with connecting blue bars (with “n.s.” in Figure 3).

We found a linear variation with similarity to cellist expertise, though this time in the ERPs. Again, we note from insets B-D of Figure 3 that the general trend for the mean ERP is a decrease from the group most similar to C expertise to the one least similar, i.e., NM. We quantified this linear increase again with the Spearman rank coefficient, finding that each ROI’s window-mean difference ERP linearly varied with similarity to C expertise (Fz: ρ = 0.85, p << 0.001, FCz: ρ = 0.95, p << 0.001, Cz: ρ = 0.87, p << 0.001).

### 3.2 Expertise Graded with Single-trial EEG Analysis

We also found gradations of expertise from a single-trial classification of the EEG data. Figure 4 shows the mean and SEM AUC values from single-trial classification of AME vs. control epochs. Color-coding indicates group membership as it did above. Each point on [−200, 1000]ms is the mean AUC value for the group at the indicated time point. Error bars indicate the SEM within that group. Particularly in the region above the Bonferroni-corrected threshold (approximately [200, 600]ms), we found a general gradation of AUC values from groups most similar to C expertise (i.e., NCSP) to those least similar (i.e., NM, NSI and S) across multiple time points.

**Figure 4.**
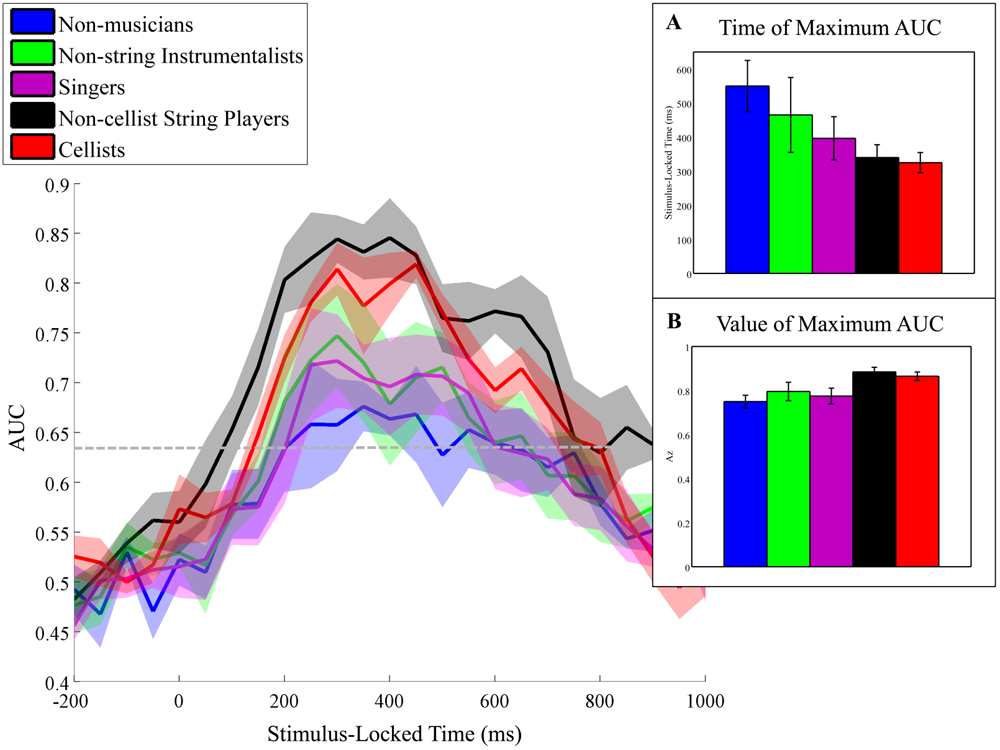
Single-trial discrimination analysis of event-related potentials as a function of subject’s mode of music production. Single-trial classification metric (AUC) vs. stimulus-locked time for each group. Dashed grey line indicates Bonferroni corrected significance line for multiple comparisons in time (p = 0.05). Group-mean values at each 50ms window are shown with standard error bars. Insets highlight group trends, such as time (A) and value (B) of maximum AUC, where means are represented by bar heights and standard errors are indicated with error bars.

Similar to the traditional ERP analysis, we performed a one-way ANOVA across groups on classification metrics and found group differences in *post hoc* single group comparisons, after correcting for multiple groups. A one-way ANOVA across groups shows a significant difference in maximum AUC value (p < 0.04, F_(4,24)_ = 3.5), though not for time of maximum AUC (p = 0.08, F_(4,24)_ = 2.4), though there appears to be a clear trend. *Post hoc* single-group comparisons of value of maximum AUC showed that NM’s value of maximum AUC is less than both that of C (p < 0.02, independent groups t-test, t(8) = −3.33) and that of NCSP (p < 0.01, independent groups t-test, t(8) = −3.81).

We also found linear variation of single-trial classification metrics with similarity to Cellist expertise. The time of maximum AUC (Figure 4A) and its value (Figure 4B) show the trend that groups more similar to C have a stronger and quicker neural response, respectively, to AMEs. We again utilized the Spearman rank coefficient finding that value of maximum AUC and its timing post-stimulus linearly varied with our assumed linear similarity scale (value of maximum AUC: ρ = 0.58, p < 0.01; timing of value of maximum AUC: ρ = −0.45, p < 0.03).

### 3.3 Single-Subject Expertise Classification from ERPs

We used an elastic net modification to the logistic regression classification technique to identify group membership from single-subject mean whole-scalp difference ERPs. In particular, we classified Cellist (C) vs. each of the other groups (NCSP, S, NSI, N) group membership at multiple post-stimulus time windows using whole-scalp ERPs as our feature space (i.e., X in Equation 2). Selecting the best classifier using the Pareto optimal criterion (see Methods and Figure S 3), the results for classifying each non-Cellist group (NCSP, S, NSI, NM) vs. the C at the indicated time windows is shown in Figure 5.

**Figure 5.**
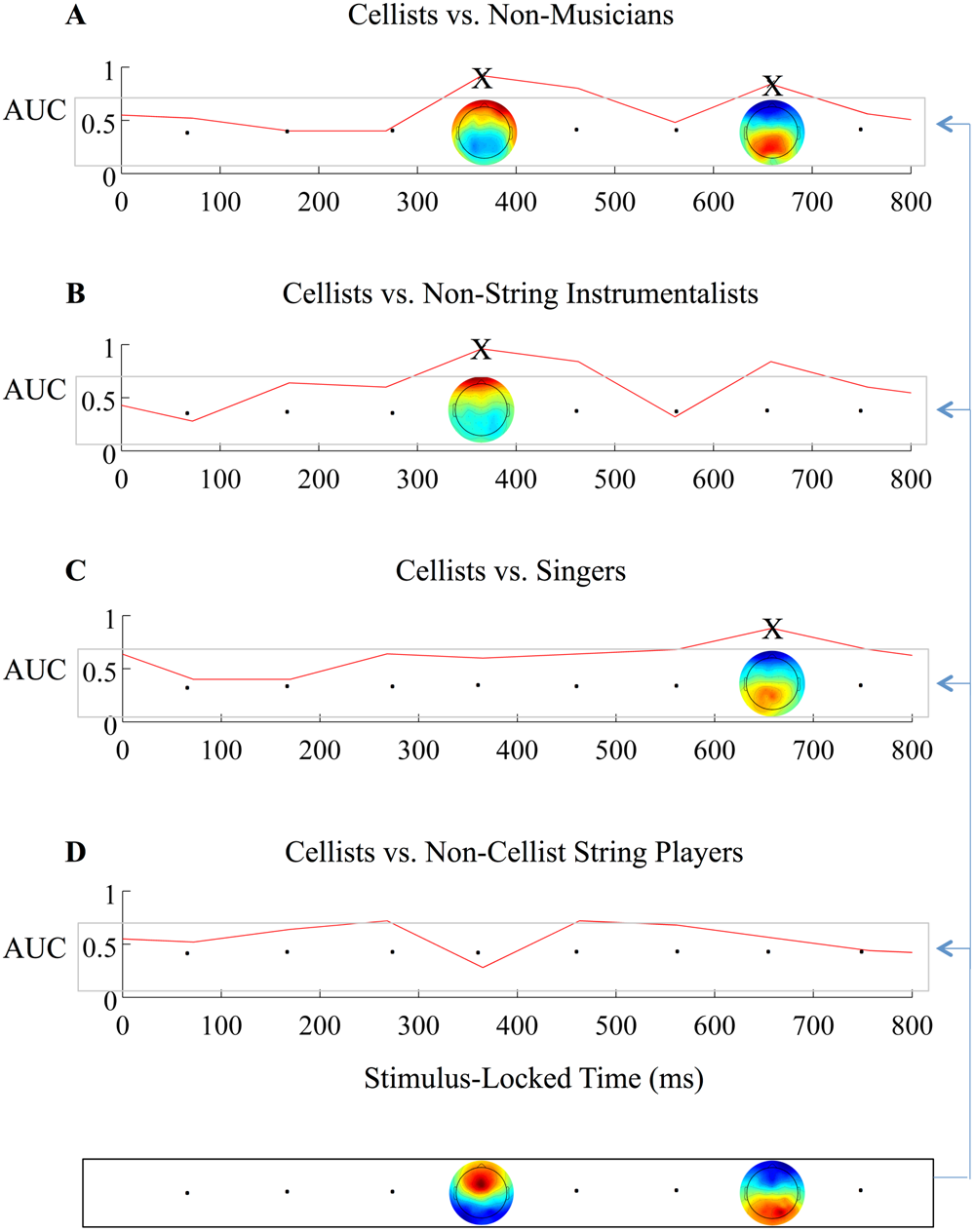
ERP classification of C vs. other groups in stimulus-locked time. Insets A-D show AUC (red line) for classification of each groups’ mean ERP in stimulus-locked time. Black ‘X’ indicates when the value of AUC is significant (p < 0.05, Bonferroni corrected for multiple comparisons in time and number of groups). Bottom of figure shows time course of mean ERPs of C. Two time points are shown because they are the superset of time points showing differences across all groups (365ms and 658ms). This time course of ERPs is classified from each other groups’ time course, specifically that of NM (A), NSI (B), S (C), and NCSP (D). Arrows indicate the four group comparisons made. From the top to the bottom of the figure, NM show differences at the P3 and N6 windows (A), the NSI show at the P3 window (B), the S show at the N6 window (C), and the NCSP show no differences (D).

The four time courses of AUC shown in Figure 5 indicate when each group showed classifiable scalp-level ERP differences on a single-subject basis from the C. As with the previous single-trial EEG epoch classification, a permutation test established the significance line shown, where there is an added Bonferroni adjustment shown in Figure 5 to account for multiple group comparisons. Time windows for which classification was significant are indicated with ‘X’ marks.

Beginning with the classification of C vs. NM, the top row (A) shows two time points that were significant (365ms and 658ms, the “early” and “late” windows respectively). The next row down (B) is a classification between the C and NSI, showing a significant classification at the early window only. The next row down (C) is a classification between the C and the S, showing a significant classification at the late window only. Finally, the last row (D) is a classification between C and NCSP, showing no significant classifications. As with previous results, we find a graded ability to classify early and late windows, proceeding from the group whose expertise is most different from that of C (top row, A) to the group whose expertise is the least different (last row, D).

Topographic ERP plots are shown at each time point for which significant classification was achieved, with the bottom box being for the C. Mirroring group expertise similarity to C, the graded change in the mean ERP plots can be seen when viewed from the top row (NM) to the last row (NCSP). A full spatio-temporal ERP topographic post-AME time course can be viewed in Supplementary Material to see these gradations more clearly (Figure S1).

### 3.4 Source Localization Reveals Motor and Executive Structures Underlie Graded Musical Expertise

We also examined group differences and gradations in expertise using current source reconstructions from the scalp EEG data. Particularly, we selected the subject-specific time window of maximum AUC and then reconstructed the sLORETA sources, comparing Cellist and non-Cellist groups’ current distributions in voxel-space. Figure 6 shows these results for comparisons of C to NM (A), NSI (B), S (C) and NCSP (D). The heat map to the right indicates t-test p-values corrected by SnPM.

**Figure 6.**
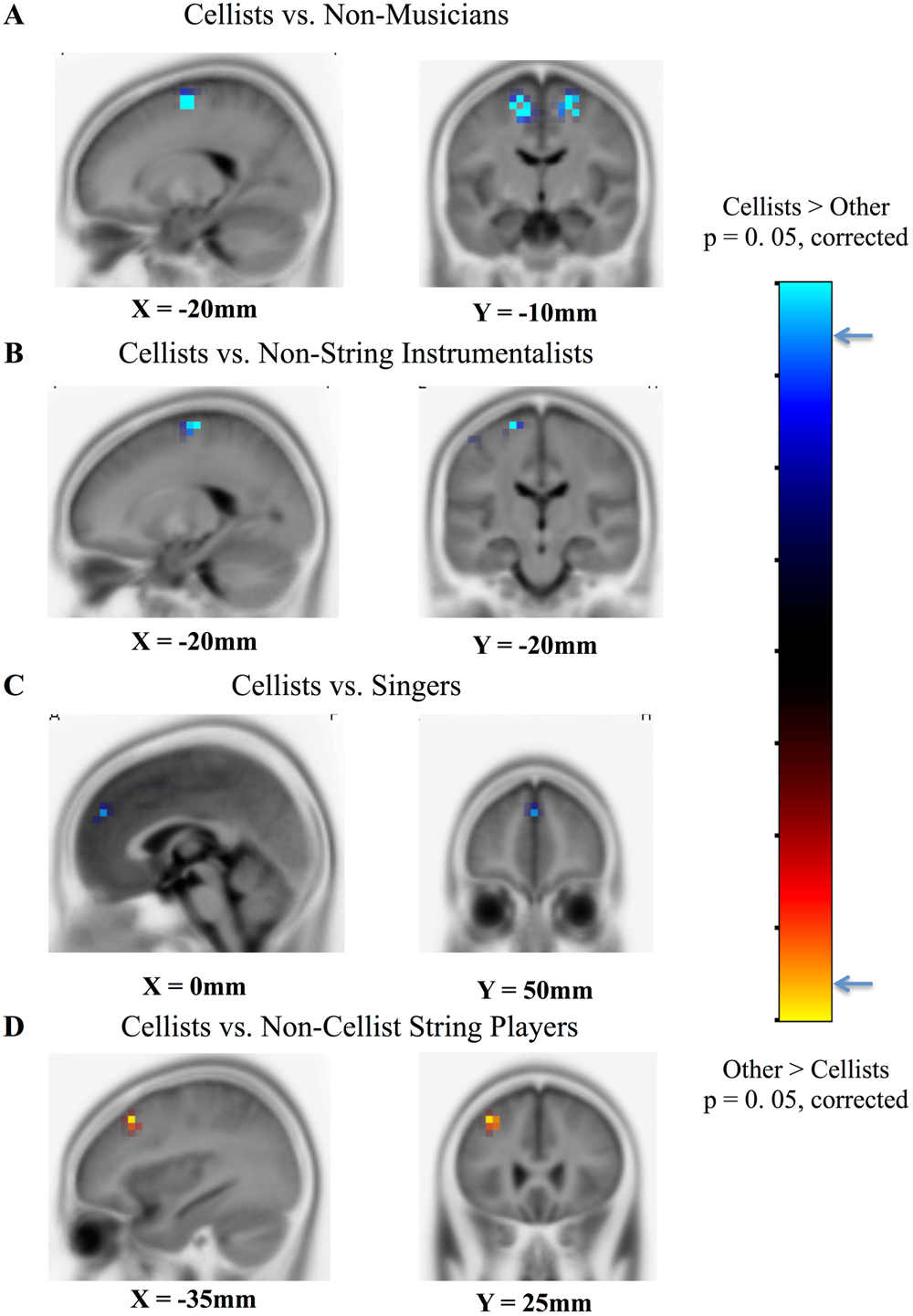
Source localization statistical maps comparing each group (NCSP, S, NSI, NM) to Cellists (C). Sagittal and coronal slices show voxels exhibiting one-sided t-test significant difference in f-ratio values, p < 0.05, Bonferroni corrected for multiple group comparisons. Heat map to the right of all insets shows where significance threshold is for each comparison. Insets A, B and C show Cellists having stronger source activity than comparison groups (NM, NSI, S), while inset D shows NCSP having stronger source activity than Cellists.

Particularly, the figure shows the t-distribution values of the log of the ratio of averages (similar to a one-way ANOVA, F(1,19)) for C > Other (purple/blue) and Other > C (orange/yellow) projected onto the cortex. Full details of which voxels showed significant differences are provided in Supplementary Material (Tables S1-S4) for each row of comparisons shown in Figure 6. All coordinates provided are MNI.

The areas most differentially activated for C compared to NM are [20, −10, 60], [−15, −10, 55] and [5, 55, 35] in the sub-gyral, medial frontal and superior frontal gyri of Brodmann Areas (BAs) 6 and 9. The areas most differentially activated for C compared to NSI are [−20, −20, 70], [−20, −15, 70] and [−20, −15, 65] in the precentral, superior frontal and middle frontal gyri of BA 6. The areas most differentially activated for C compared to S are [0, 50, 30] and [−50, 20, 40] in the medial frontal and cingulate gyri of BAs 9 and 32. Finally, the areas most differentially activated for NCSP compared to C contain [−35, 25, 45] and [−30, 20, 40] in the middle frontal and precentral gyri of BAs 8 and 9.

As reported in our earlier work (Sherwin and Sajda, 2013), we found that the brain region of peak activity for C vs. NM has also been implicated in music imagery tasks in pianists. Also, we also found strong right lateralized activity for C vs. NM when we consider SnPM-corrected voxels out to p < 0.05 (31 left voxels vs. 50 right voxels, 17 of which are in right frontal cortices). These results hold also for C vs. NSI (p < 0.05, 34 left voxels vs. 0 right voxels), where it is important to note that the laterality is completely reversed. This is likely because the NSI used are right-handed trumpeters, who modify pitch with the valves played by the right hand; C use both hands to play a given pitch, but the left hand serves as the predominant guide for pitch adjustment (i.e., the “fret” hand). We found slight right lateralization of activity for C vs. S (p < 0.05, 13 left voxels vs. 16 right voxels), but not centered on the motor cortices; rather, the source of differential activity is focused at the cingulate cortex, which is used for error monitoring (Lavin et al., 2013; Shenhav et al., 2013). Finally, we find strong left lateralized activity for C vs. NCSP in the frontal cortices (p < 0.05, 6 left voxels vs. 0 right voxels), though there is no difference in motor areas between these groups. Generally, the progression from more disparate expertise groups (A, i.e., NM vs. C) to more similar groups (D, i.e., NCSP vs. C) shows a difference in source activity moving from motor areas towards executive frontal areas.

## 4 Discussion

The objective of this research was to investigate and identify potential neural correlates differentiating musicians based on their modes of musical production. Importantly, this study was done in terms of how these correlates are expressed when subject’s listen to a musical piece absent their active participation in its generation. Specifically subjects to count anomalous musical events in a cello piece and report at the end of the experiment the number of anomalies they detected. We recorded EEG during the task and analyzed this data as a function of the subjects’ mode of musical production. Our analysis revealed a general gradation in EEG correlates, both ERPs and single-trial signatures, as a function of the deviation of the mode of production from the cellist. Furthermore, we localized these signatures to motor and executive-function cortical areas that could account for this differentiation in the mode of music production. We contextualize these finding in the relevant literature below.

### 4.1 Traditional ERP and Behavioral Analysis Reveals Graded Musical Expertise

Traditional ERP and behavioral analyses comprise the foundation of EEG research into musical expertise (Koelsch et al., 2000; Koelsch et al., 2002b; Koelsch et al., 1999; Tervaniemi et al., 2006; Tervaniemi et al., 2005). Using this classic methodology, we found that the level of “cellist expertise”, as defined by the musician’s distance to the cellist in terms of their mode of musical production, was generally tracked by trial-and window-averaged ERP values (Figure 3). CFz and Cz electrodes showed consistent trends in their mean values ERP differences, while electrode Fz was consistent except for the non-cellist string players (NCSPs). Across the group, we found a significant correlation in terms of the mean ERPs and mode of music production relative to cellist. This correlation was also seen in the behavioral accuracy. Interestingly, our source localization findings show that the NCSPs and Cs differ primarily in frontal cortical activity and not motor areas, as do cellists and the other groups. Further research would be needed to fully investigate why executive control centers are differentiating in this case.

### 4.2 Single-trial ERP Analysis Reveals Graded Musical Expertise in Completely Data-driven Fashion

The overwhelming benefit provided by our single-trial EEG analysis is that signal-to-noise can be increased by integrating across the sensors and differences between groups can be identified not just in terms of the means of the activity but its variation, on a trial-to-trial basis, relative to evoked activity generated by the AME. Our single-trial analysis (Figure 4) is consistent with our ERP finding (Figure 4) showing the same trends across the groups. Importantly, the single-trial results also enable us to identify when during the trial the AME is expressed in the underlying neural activity.

With these single-trial results in hand, we used the timing and value of maximum discrimination, as indexed by the AUC of the classifier, to correlate this to the mode of music production relative to cellist. We found trends that demonstrate a graded relationship, which previously was only seen as a binary differentiation between NM to musicians (Musacchia et al., 2008; Sherwin and Sajda, 2013) and other expert to novice groups (Muraskin et al., 2016; Muraskin et al., 2015). In particular, we find that the more similarity in musical sound production between the subject and the stimulus, the faster the neural response of the subject to the AME in that stimulus. We were not able to observe this when using the trial averaged ERPs at Fz and CFz.

### 4.3 Single-subject Classification from ERP Data Demonstrates Predictive Ability to Grade Musical Expertise Type

Just as it had been used to learn discriminating features of AME epochs, we also used machine learning to naïvely learn features of expertise (see Figure 2). By employing necessary thresholds for multiple comparisons, we find a decomposition in Figure 5 of what sensors and when their response drove expertise classification.

Beginning with NMs vs. Cs, we see that two windows provide suitable data to discriminate these two subject groups, particularly the windows of the P300 and the P600 (Figure 5A). Previous research has shown that the former of these windows drives expectancy violation for general stimuli (Goldman et al., 2009; Walz et al., 2013; Walz et al., 2014), while the latter window drives semantic content expectancy violation (Beim Graben et al., 2008; Friederici, 2002). When we do the same classification for NSIs vs. Cs, we only see that the P300 window can be used to classify significantly (Figure 5B). Continuing onto the classification of Ss vs. Cs, we only now see that the P600 window can be used for significant classification (Figure 5C). Finally, doing the classification between NCSPs and Cs, we find that neither of these windows (nor any others) can be used to significantly classify.

This decomposition of expertise classification is completely data driven. Progressing from the groups with last similarity in cellist sound production expertise (Figure 5A), to the groups with the most (Figure 5D), we find that the P300 and P600 windows are the compositional parts that drive the hypothesized trends seen in traditional/single-trial EEG and behavioral metrics. Figure 5 though is a trend in time and across groups. Furthermore, it is driven by all of the sensor measurements and all of the time points in conjunction with naïve classification, rather than *a priori* hypotheses about where or when to look for markers of expertise.

### 4.4 Auditory-motor Connections Drive Musical Expertise Similarities and Differences

The final piece to decomposing the hypothesized trends across increasingly similar groups via their expertise comes form the source localization results (Figure 6, full MNI and structure details in Table S2,Table S3,Table S4, Table S5). Here, we show that the two groups with the least similarity to Cs (i.e., NMs and then NSIs) differ the most at maximum discrimination of AME epochs in the motor cortical areas. And the two groups with the most similarity to Cs (i.e., Ss and then NCSPs) differ the most at maximum discrimination of AME epochs in frontal (i.e., executive) cortical areas – no differences were found in motor areas for these comparisons.

As hypothesized in earlier work (Sherwin and Sajda, 2013), this heightened use of motor cortical areas in Cs vs. NMs is believed to be due to the mechanism by which Cs would modify produced pitches on the cello with their bow-and fret-hands, where the latter plays a more dominant role in pitch modification. As reported earlier, we found differentially more activity on the fret-hand side among Cs (i.e., the right side of the brain for left fret-hands, i.e., “right-handed” cellists). Now, we confirm the veracity of this hypothesis by adding the evidence provided by NSIs, particularly trumpeters. We find that the source response in cortical motor areas of trumpeters is stronger on the left side of the brain, i.e., for right-handed trumpeters who play valves with their right-hand (see Table S1 for handedness details). Importantly, the source localization results comparing Cs to both NCSPs and Ss provide a valuable control test to this hypothesis. We find no significant difference in motor cortical areas from the SnPM method of assessing differential activity in sLORETA estimates (Figure 6), rather finding that frontal cortices show significant differences. A possible future direction of this research would be to identify whether this result is explained by the fact that Cs, NCSPs and Ss are non-fixed-pitch modes of music production, compared to NSIs, which generally are.

## 5 Supplementary Material

### 5.1 Detailed Block Design Implementation

The 8 blocks of five music excerpts unfolded in a pseudorandomized order to minimize the possibility of subject habituation to the timing of AMEs. An example implementation is shown in Figure S 1, with the inset showing Figure 1.

**Figure S 1.**
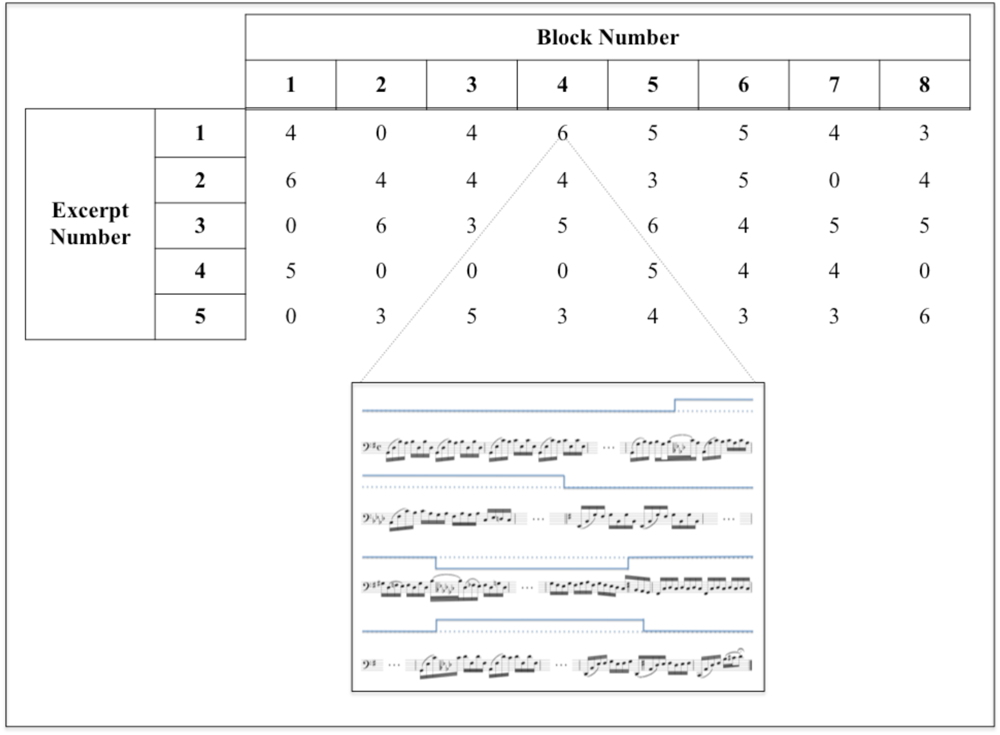
(Reprinted from (Sherwin and Sajda, 2013)) Example of our randomized block design. Within each block, the ordering of excerpt playback was pseudorandomized. Blocks contained anywhere from zero to two control excerpts in which no key-changes occurred. The inset shows an example key-change excerpt for which six key-changes occurred. Although the shown inset begins and ends in the original recording key (G), this was not always the case, so as not to bias the detection simply being in the non-original recording key.

### 5.2 Detailed Background Information on Subjects

All subjects were recruited based on criterion of general musical and voice/instrument training. With the exception of one Non-Musician, all subjects were right-handed. Full details for all twenty-seven subjects are shown in Table S1.

**Table S1.** Full background detail of all subjects. Each column shows from left to right the unique subject identifier, the age, the age of musical training start, the age of voice or instrument training start, and handedness. For the subject identifier, the leading letters correspond to Cellists (C), Non-Cellist String Players (NCSP), Singers (S), Non-String Instrumentalists (NSI) and Non-professional Musicians (NM). Handedness is indicated by right-handed (R) or left-handed (L).

|  | Subject Identifier | Subject Age (years) | Age of Musical Training Start (years) | Age of Voice/Instrument Training Start (years) | Handedness |
| --- | --- | --- | --- | --- | --- |
| Cellists | C1 | 51 | 6 | 10 | R |
|  | C2 | 39 | 4 | 7 | R |
|  | C3 | 28 | 8 | 8 | R |
|  | C4 | 53 | 9 | 9 | R |
|  | C5 | 35 | 5 | 9 | R |
| Non-Cellist String Players | NCSP1 | 28 | 6 | 6 | R |
|  | NCSP2 | 29 | 6 | 6 | R |
|  | NCSP3 | 40 | 8 | 8 | R |
|  | NCSP4 | 40 | 4 | 4 | R |
|  | NCSP5 | 29 | 10 | 10 | R |
| Singers | S1 | 28 | 8 | 14 | R |
|  | S2 | 24 | 5 | 5 | R |
|  | S3 | 46 | 3 | 3 | R |
|  | S4 | 54 | 4 | 4 | R |
|  | S5 | 44 | 3 | 6 | R |
|  | S6 | 29 | 7 | 12 | R |
|  | S7 | 32 | 8 | 8 | R |
| Non-String Instrumentalists | NSI1 | 30 | 6 | 10 | R |
|  | NSI2 | 39 | 5 | 10 | R |
|  | NSI3 | 19 | 5 | 5 | R |
|  | NSI4 | 21 | 5 | 9 | R |
|  | NSI5 | 25 | 6 | 9 | R |
| Non-Musicians | NM1 | 27 | 8 | 27 | R |
|  | NM2 | 41 | 23 | 41 | R |
|  | NM3 | 28 | 23 | 28 | L |
|  | NM4 | 28 | 8 | 8 | R |
|  | NM5 | 30 | 30 | 30 | R |

### 5.3 Additional Traditional ERP Analyses

We complemented the classification results on mean difference ERPs that allowed us to identify subjects’ group memberships. A full spatio-temporal topographic scalp plot shows the progression in post-stimulus time windows of the mean difference ERP at the scalp.

We found that early activity (100ms) was not spatially coherent, except in Non-Cellist String Players (NCSP). This result corroborates the early significant discrimination we see from this group in Figure 4. By 300ms and 400ms, all groups show coherent frontal positivity and occipital negativity. This corroborates also the trends seen in Figure 4, where all groups show significant discrimination by these time points. It is important to note that the musician groups (i.e., all but NM) show a left lateralization in the frontal positivity. As all musician subjects were right-handed (see Table S1), this is likely due to the motor component of AME detection, a result corroborated by the sLORETA findings. Finally, the frontal positivity of the C is the most focused and occipital of the five groups, placing it closest to motor cortical areas. As with the laterality results, this corroborates also the greater sLORETA activity seen in this group compared to the NM and, in fact, the progression from NM to C in Figure S 2 tracks those source results well.

**Figure S 2.**
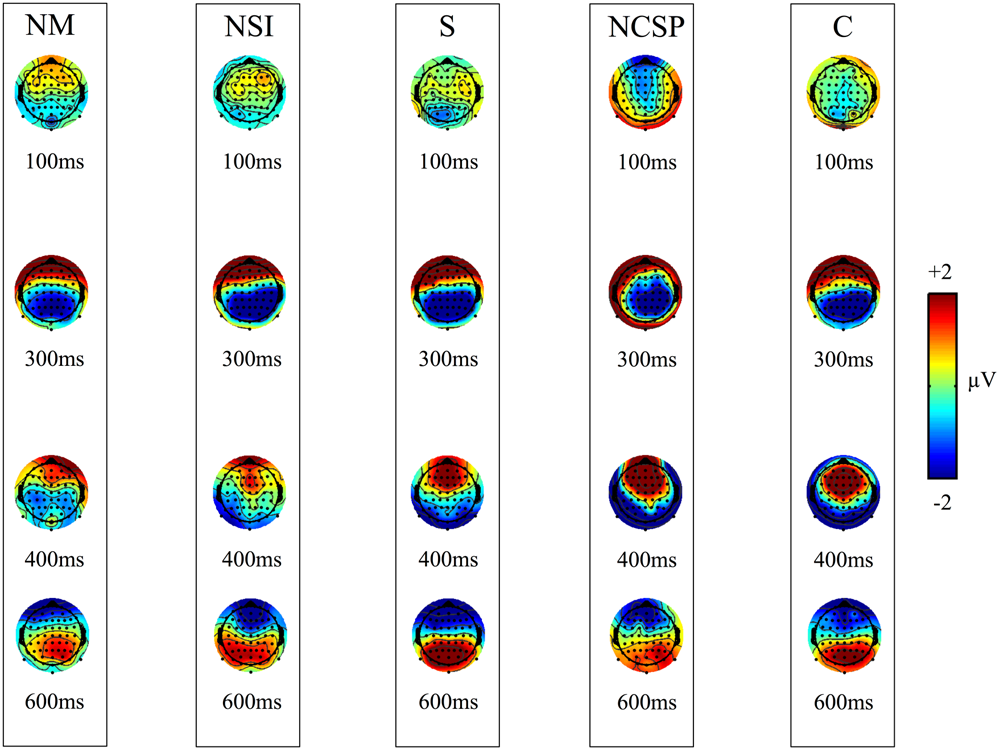
Time course of post-stimulus topographic ERP scalp plots. Each column shows topographic mean ERP difference plots between AME and control conditions at the indicated time. Column abbreviations for the five groups are as follows: NM = Non-professional Musicians, NSI = Non-String Instrumentalists, S = Singers, NCSP = Non-Cellist String Players, C = Cellists. All plots are color-coded to voltage according to the color bar at right. For each column, the general pattern in time is that early activity is not coherently spatially localized (e.g., at 100ms). But from 300ms to 600ms, the activity moves from a spatially coherent frontal positivity with occipital negativity to the reverse. Differences at each indicated time point across groups are visually clear. Of particular note is at 300ms, when musicians’ positivity is left lateralized, whereas the NM are not.

### 5.4 Details on Elastic Net Regularized Logistic Regression

The single-subject classification of expertise relied on an elastic net regularized logistic regression. In this technique, the AUC of a non-elastic net approach is replaced with the optimum value along the Pareto frontier defined between AUC and mean probability of feature selection. This selection is illustrated in Figure S 3 for the example of the 368ms time point of the classification between C and NM shown in Figure 5A.

**Figure S 3.**
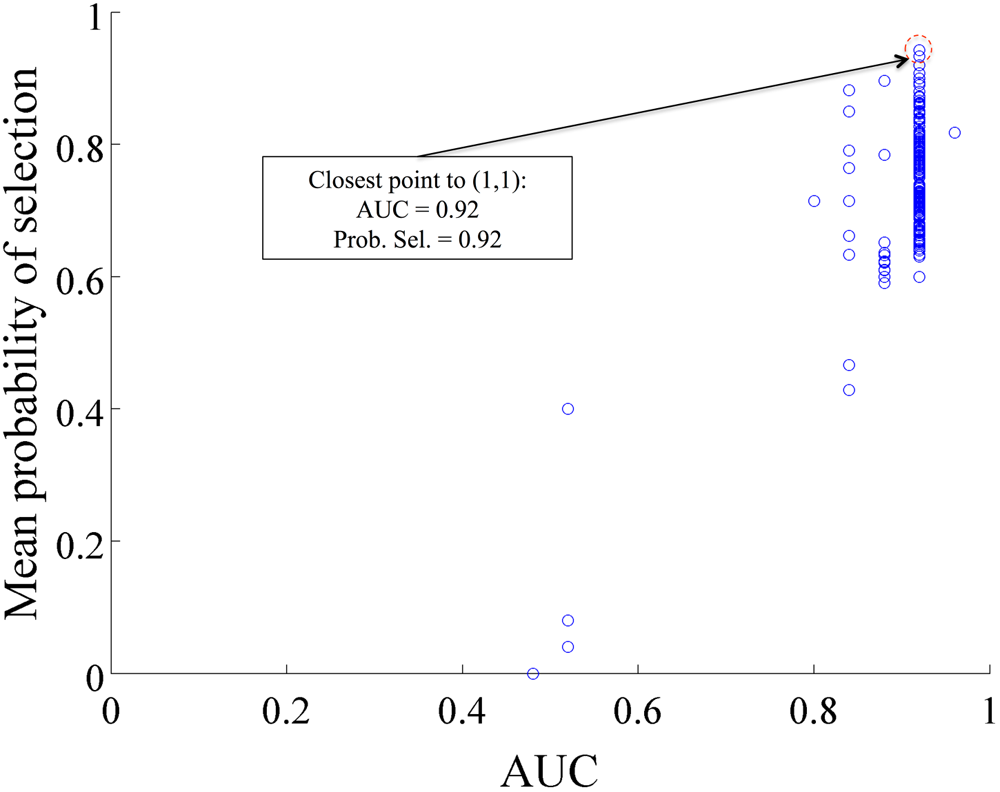
Example Pareto Frontier for AUC selection. Each point (blue circle) represents a solution in the regularized logistic regression elastic net space. The point closest to AUC = 1.0 and Mean probability of selection = 1.0 is the optimum point along the Pareto frontier, having the optimum tradeoff between high AUC and consistent selection of the same features across the 10-fold leave-one-subject-out cross validation.

### 5.5 Detailed Source Localization Results and Parameters

We found multiple voxels showing significant differences between the Cellist and other groups. These voxels tended to be clustered in certain regions of the cortex discussed in the Results and Discussion sections of the paper. Full location details in MNI coordinates with accompanying neural structure information are provided here for C vs. NM (Table S2, reprinted from (Sherwin and Sajda, 2013)), C vs. NSI (Table S3), C vs. S (Table S4) and C vs. NCSP (Table S5).

**Table S2.** MNI coordinates, Brodmann areas and cortical structures showing greater neuronal source activity among C than NM at peak discrimination between key-changes and controls. After correcting for multiple comparisons using SnPM, the 15 points shown here are the only voxels at which C’ neuronal sources are greater than those of NM (p<0.01, independent groups t-test).

| X(MNI) | Y(MNI) | Z(MNI) | Voxel t-value | Brodmann area | Structure |
| --- | --- | --- | --- | --- | --- |
| 20 | -10 | 60 | -3.102 | 6 | Sub-Gyral |
| -15 | -10 | 55 | -3.091 | 6 | Medial Frontal Gyrus |
| 5 | 55 | 35 | -3.079 | 9 | Superior Frontal Gyrus |
| 10 | 55 | 35 | -3.068 | 9 | Superior Frontal Gyrus |
| -15 | -15 | 55 | -3.044 | 6 | Medial Frontal Gyrus |
| 20 | -5 | 60 | -3.008 | 6 | Sub-Gyral |
| 5 | 50 | 35 | -2.999 | 9 | Superior Frontal Gyrus |
| 20 | -5 | 65 | -2.969 | 6 | Middle Frontal Gyrus |
| 25 | -5 | 60 | -2.968 | 6 | Sub-Gyral |
| 20 | -10 | 65 | -2.963 | 6 | Middle Frontal Gyrus |
| -15 | -15 | 60 | -2.960 | 6 | Medial Frontal Gyrus |
| 25 | -5 | 65 | -2.941 | 6 | Middle Frontal Gyrus |
| -15 | -10 | 65 | -2.939 | 6 | Middle Frontal Gyrus |
| -10 | -10 | 55 | -2.933 | 6 | Medial Frontal Gyrus |
| -20 | -10 | 60 | -2.902 | 6 | Sub-Gyral |

**Table S3.** MNI coordinates, Brodmann areas and cortical structures showing greater neuronal source activity among C than NSI at peak discrimination between key-changes and controls. After correcting for multiple comparisons using SnPM, the 10 points shown here are the only voxels at which C’ neuronal sources are greater than those of NSI (p<0.03, independent groups t-test).

| X(MNI) | Y(MNI) | Z(MNI) | Voxel t-value | Brodmann area | Structure |
| --- | --- | --- | --- | --- | --- |
| -20 | -20 | 70 | -2.836 | 6 | Precentral Gyrus |
| -20 | -15 | 70 | -2.771 | 6 | Superior Frontal Gyrus |
| -20 | -15 | 65 | -2.727 | 6 | Middle Frontal Gyrus |
| -15 | -15 | 70 | -2.690 | 6 | Superior Frontal Gyrus |
| -15 | -20 | 70 | -2.679 | 6 | Precentral Gyrus |
| -15 | -10 | 70 | -2.657 | 6 | Superior Frontal Gyrus |
| -15 | -15 | 60 | -2.650 | 6 | Medial Frontal Gyrus |
| -20 | -10 | 70 | -2.635 | 6 | Superior Frontal Gyrus |
| -15 | -10 | 65 | -2.632 | 6 | Middle Frontal Gyrus |
| -25 | -15 | 70 | -2.61998 | 6 | Precentral Gyrus |

**Table S4.**
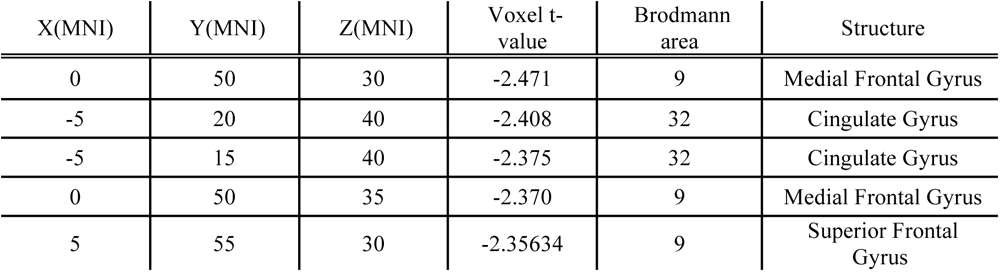
MNI coordinates, Brodmann areas and cortical structures showing greater neuronal source activity among C than S at peak discrimination between key-changes and controls. After correcting for multiple comparisons using SnPM, the 5 points shown here are the only voxels at which C’ neuronal sources are greater than those of S (p<0.03, independent groups t-test).

**Table S5.** MNI coordinates, Brodmann areas and cortical structures showing greater neuronal source activity among NCSP than C at peak discrimination between key-changes and controls. After correcting for multiple comparisons using SnPM, the 6 points shown here are the only voxels at which NCSP’ neuronal sources are greater than those of C (p<0.03, independent groups t-test).

| X(MNI) | Y(MNI) | Z(MNI) | Voxel t-value | Brodmann area | Structure |
| --- | --- | --- | --- | --- | --- |
| -35 | 25 | 45 | 3.178 | 8 | Middle Frontal Gyrus |
| -30 | 20 | 45 | 3.170 | 8 | Middle Frontal Gyrus |
| -30 | 25 | 45 | 3.137 | 8 | Middle Frontal Gyrus |
| -30 | 25 | 40 | 3.119 | 9 | Middle Frontal Gyrus |
| -30 | 20 | 40 | 3.116 | 9 | Precentral Gyrus |
| -35 | 25 | 40 | 3.10197 | 9 | Middle Frontal Gyrus |

**Table S6.** Parameters used in the sLORETA statistical analysis. With 10 samples per subject group, the maximum number of permutations for SnPM is 2^10 and so we exceeded that in our analysis. Our statistical analysis used the log of ratio of averages and an independent groups t-test.

| **sLORETA Parameters** |  |
| --- | --- |
| Name | Value |
| No normalization | TRUE |
| Independent groups, test A=B | TRUE |
| No baseline | TRUE |
| All tests for all Time-Frames/Frequencies | TRUE |
| Log of ratio of averages (similar to log of f-ratio) | TRUE |
| Number of randomizations | 5000 |

## 6. Acknowledgements

This research was sponsored by the Army Research Laboratory and was accomplished under Cooperative Agreement Number W911NF-15-2-0074. Additionally, this work was supported by grants from the Army Research Office (W911NF-11-1-0219), the National Institutes of Health (R01-MH085092), the Army Research Laboratory under Cooperative Agreement Number W911NF-10-2-0022 and in part by an appointment to the U.S. Army Research Laboratory Postdoctoral Fellowship Program administered by the Oak Ridge Associated Universities through a contract with the U.S. Army Research Laboratory. The views and conclusions contained in this document are those of the authors and should not be interpreted as representing the official policies, either expressed or implied, of the Army Research Laboratory of the US Government. The US Government is authorized to reproduce and distribute reprints for Government purposes notwithstanding any copyright notation herein.

